# Sigma70Pred: A highly accurate method for predicting sigma70 promoter in prokaryotic genome

**DOI:** 10.1101/2021.06.29.450448

**Authors:** Sumeet Patiyal, Nitindeep Singh, Mohd. Zartab Ali, Dhawal Singh Pundir, Gajendra P.S. Raghava

## Abstract

Sigma70 factor plays a crucial role in prokaryotes and regulates the transcription of most of the housekeeping genes. One of the major challenges is to predict the sigma70 promoter or sigma70 factor binding site with high precision. In this study, we trained and evaluate our models on a dataset consists of 741 sigma70 promoters and 1400 non-promoters. We have generated a wide range of features around 8000 which includes Dinucleotide Auto-Correlation, Dinucleotide Cross-Correlation, Dinucleotide Auto Cross-Correlation, Moran Auto-Correlation, Normalized Moreau-Broto Auto-Correlation, Parallel Correlation Pseudo Tri-Nucleotide Composition, etc. Our SVM based model achieved maximum accuracy 97.38% with AUROC 0.99 on training dataset, using 200 selected features. In order to check the robustness of our model we have tested our model on the independent dataset made by using latest version of RegulonDB10.8, which included 1134 sigma70 and 638 non-sigma70 promoters, and able to achieve accuracy of 90.41% with AUROC of 0.95. We have developed a method, Sigma70Pred, which is available as webserver and standalone packages at https://webs.iiitd.edu.in/raghava/sigma70pred/. The services are freely accessible.

## Introduction

Promoters and enhancers regulate the fate of a cell and the expression of genes. Promoters are generally located upstream of genes’ transcription start sites (TSS) responsible for switching on or off the respective gene. In prokaryotes, promoters are recognized by the holoenzyme, which is made up of RNA polymerase and a related sigma factor. There are various types of sigma factors responsible for different functions, such as sigma54 controls the transcription of genes responsible for the modulation of cellular nitrogen levels, sigma38 controls stationary phase genes, sigma32 regulates heat-shock genes, and sigma24 and sigma18 controls the extra-cytoplasmic functions [1]. The number associated with each sigma factor represents the molecular weight. Sigma70 factor is a crucial factor as it regulated the transcription of most of the housekeeping genes. Sigma70 promoter comprises two well-defined short sequences located at -10 and -35 base pairs upstream of TSS, known as pribnow box and -35 region [2]. It is essential to classify the promoters in a genome as it can aid in illuminating the genome’s regulatory mechanism and disease-causing variants within cis-regulatory elements. The area of promoters is of great interest as people pay great attention to their importance not only in developmental gene expression but also in environmental response. Due to the advancement in sequencing technology, the data is growing exponentially, and hence the classification of the promoter region is a crucial problem because the standard procedures are expensive in terms of time, and performance [3,4].

In the past, ample of methods have been developed for predicting sigma70 promoters. IMPD [5], is based on increment of diversity, which achieved an accuracy of 87.9%. This method was trained on RegulonDB [6] dataset that contains 741 E. coli sigma70 promoters. Z-curve-based approach [7] attains the maximum accuracy of 96.1% by using a smaller dataset that comprises 576 sigma70 promoters and 825 non-sigma70 promoters. PseZNC [8] is based on a multi-window Z-curve approach and gained the maximum accuracy of 84.5% using the dataset from RegulonDB9.0 [6]. 70Propred [9] has incorporated features like position-specific trinucleotide propensity based on single-stranded characteristic (PSTNPss) and electron-ion potential values for trinucleotides (PseEIIP), and reported the 95.56% accuracy.

In the present study, we have developed a computational method called as Sigma70Pred, to classify the sequences in sigma70 promoter and non-promoter. In this study, dataset used for benchmarking is same as used in 70Propred. One of the objectives of this study is to improve the prediction performance of models on a large and recent dataset. A web server and python and docker-based standalone software have been developed to serve the scientific community for predicting the sigma70 promoters.

## Materials And Methods

### Dataset generation

In order to train and test our models using cross-validation, we obtained training dataset from RegulonDB9.0 [6]. It contains 741 sigma70 promoters and 1400 non-promoters, and each sequence is of length 81. The same data has been used previously by 70Propred. In order to validate our model on external dataset or independent dataset, we have extracted the dataset from RegulonDB 10.8, which comprises 1134 sigma70 and 638 non-sigma70 promoters. There is no identical sequence in training and independent dataset. The datasets can be downloaded from our server.

### Overall workflow

The comprehensive workflow for Sigma70Pred is shown in Figure 1.

**Figure 1:**
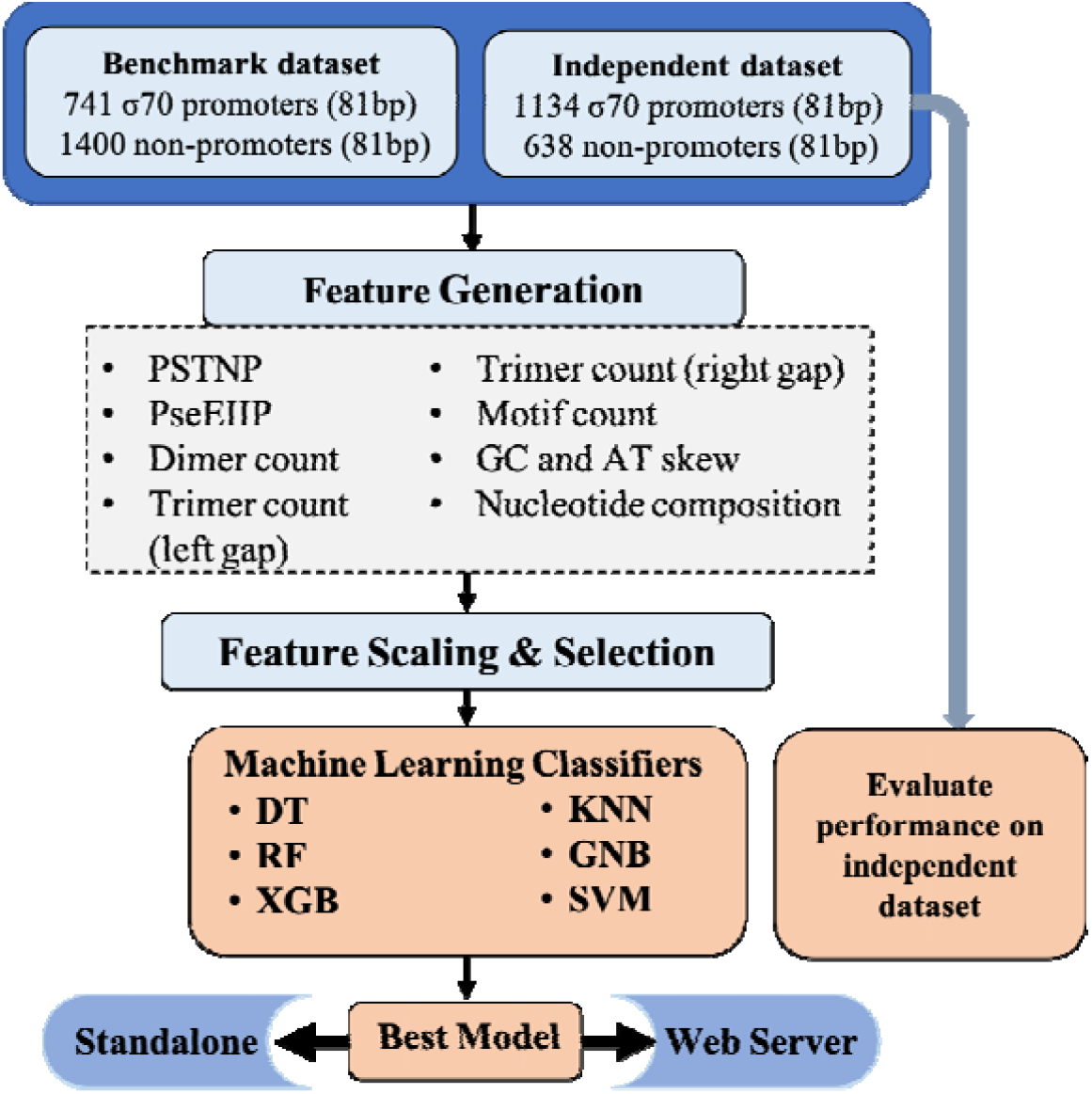
Architecture of Sigma70Pred.

### Feature generation

We have generated wide range of features like Position-Specific Tri-Nucleotide Propensity (PSTNPP), Electron-Ion Interaction Pseudopotentials of trinucleotide (EIIIP) [9], dimer count, trimer count, motif counts, GC and AT skew [10], Dinucleotide Auto-Correlation (DAC), Dinucleotide Cross-Correlation (DCC), Dinucleotide Auto Cross-Correlation (DACC) [11], Moran Auto-Correlation (MAC), Normalized Moreau-Broto Auto-Correlation (NMBAC) [12], and Parallel Correlation Pseudo Tri-Nucleotide Composition(PC_PTNC) [13], which resulted in 8465 features. Then, we have used the Min-Max scaler from the scikit-learn library [14] to scale down the values of the feature we have constructed. This is done in order to bring the values of features to the same range (usually 0 and 1). Then we have implemented Recursive Feature Elimination (RFE) [14] for the feature selection. In this technique, we eliminate some of the features from our dataset and construct the model using the base estimator function on a new set of features, and performance is evaluated on the basis of certain metrics. This process is repeated until we are left with the desired number of features. Details of each feature and processing of features are explained in the Supplementary file.

### Model development

In this study, we developed models for predicting sigma70 promoters using wide range of machine learning techniques such as decision tree (DT), random forest (RF), k-nearest neighbor (KNN), extreme gradient boosting (XGB), gaussian naïve bayes (GNB), and support vector machine (SVM) [14]. We got the best performance using SVM based model. Our best model on training dataset was evaluated on independent dataset (obtained from RegulonDB 10.8).

### Five-fold cross-validation

In order to avoid the biasness and test the prediction models’ performance, we have implemented five-fold cross-validation. In this approach, the complete dataset is divided into five parts, the model is trained on four out of five parts, whereas the model is tested on the left part, and the performance is recorded. The same process is iterated five times so that each part gets the chance to be used for the purpose of testing. The overall performance is calculated by taking the mean of all five iterations [15].

### Measures of performance

To assess the performance of generated prediction models, we have used various threshold-dependent and independent parameters. We have considered sensitivity that is, percent of sigma70 samples classified correctly; specificity that is, percent of non-sigma70 samples classified as negative; accuracy that is, percentage of samples which are correctly predicted by the model; and Matthews correlation coefficient (MCC) that explains the relationship between the observed and predicted value, under threshold-dependent parameters, whereas, in threshold-independent measures, we have considered Area Under the Receiver Operating Characteristics (AUROC) which is the relation between true positive rate and false positive rate. The AUROC was computed and plotted using the pROC package [16] of R. The equations depicting the threshold-dependent parameters are as follows:

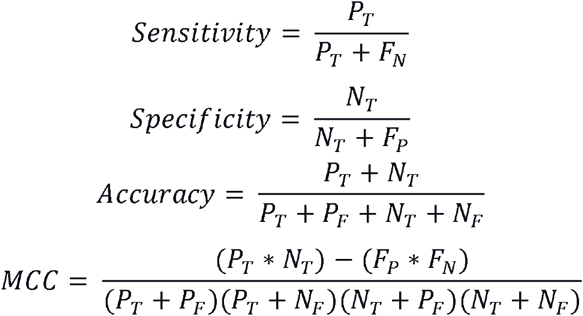

where, P_T_ refers to number of true positives; P_F_ refers to number of false positives; N_T_ refers to number of true negatives; and N_F_ refers to number of false negatives.

## RESULTS

### Compositional analysis

In order to assess the proportion of the nucleic acids in the sigma70 promoter and non-promoter, we have calculated the mono-nucleotide composition. As shown in Figure 2, nucleic acid adenine and thymine are abundant in sigma70 promoter sequences, whereas cytosine and guanine are higher in percentage in the case of non-promoter sequences.

**Figure 2:**
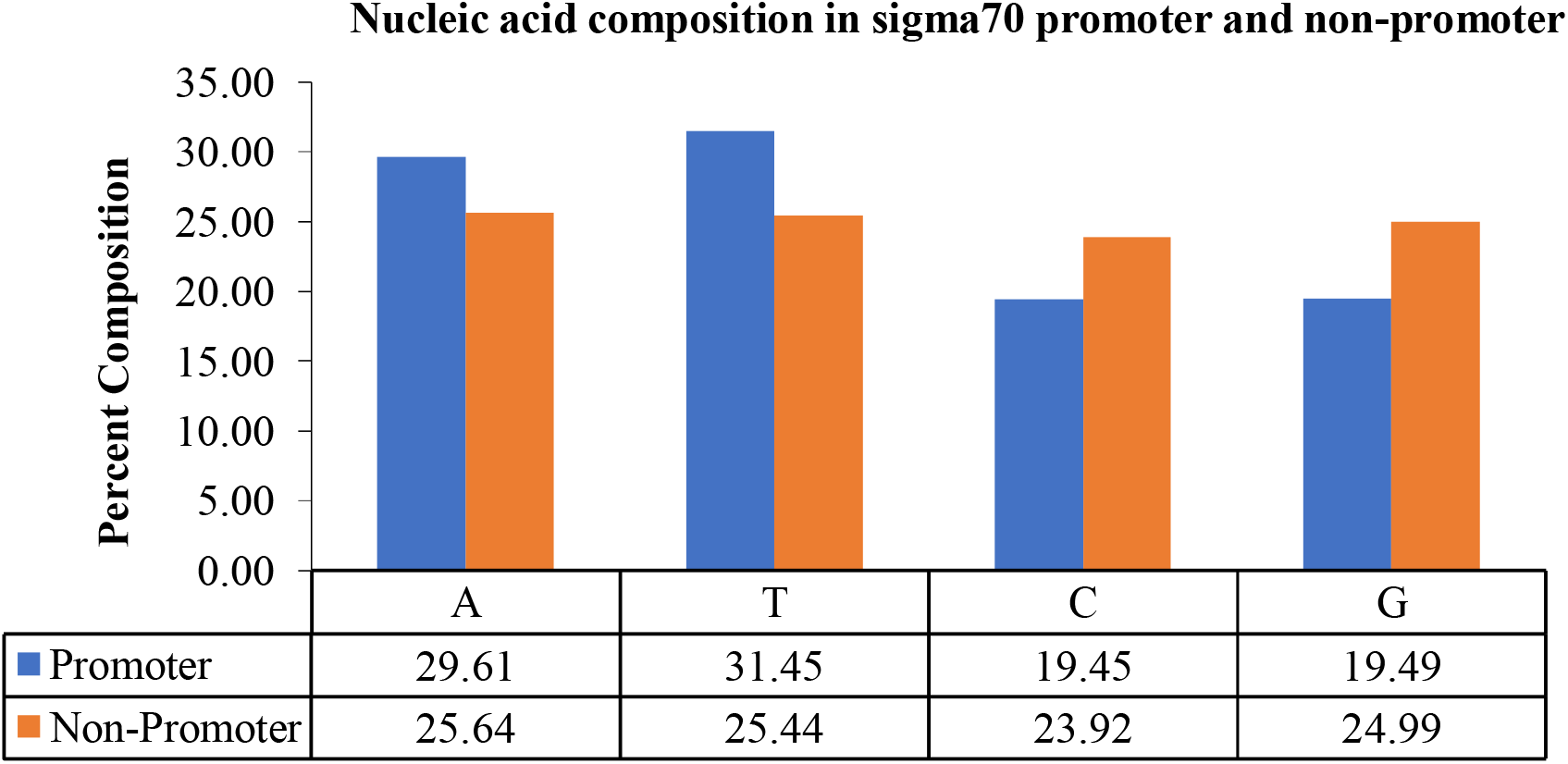
Mono-nucleotide composition of sigma70 promoters and non-promoters.

### Performance of machine learning classifiers on benchmark dataset

Initially, we have generated around 8000 nucleotide based features, and then selected 200 most relevant features using feature scaling method min-max scaler and feature selection method RFE. Using these selected features, we have generated various models by implementing various machine learning techniques. The SVM based model performed best among all the other classifiers with 97.38% accuracy, 0.99 AUROC, and 0.94 MCC on benchmark dataset as shown in Table 1.

**Table 1:** Performance of various machine learning classifiers on benchmark dataset.

| Classifier | Sensitivity | Specificity | Accuracy | AUROC | MCC |
| --- | --- | --- | --- | --- | --- |
| DT | 74.49 | 87.14 | 82.77 | 0.808 | 0.62 |
| RF | 92.04 | 91.57 | 91.73 | 0.977 | 0.82 |
| XGB | 91.90 | 92.14 | 92.06 | 0.980 | 0.83 |
| KNN | 90.15 | 91.79 | 91.22 | 0.958 | 0.81 |
| GNB | 88.66 | 88.71 | 88.70 | 0.955 | 0.76 |
| **SVM** | **97.44** | **97.36** | **97.38** | **0.996** | **0.94** |

### Performance comparison with existing methods

There are ample of methods which are trained and evaluated on the same benchmark dataset such as, 70ProPred [9], iPro70-FMWin [10], PseZNC [8], Z-Curve [7], and IPMD [5]. We have compared the performance of Sigma70Pred with existing prediction methods for sigma70 promoters prediction and found out that Sigma70Pred has outperformed all the existing methods, as shown in Table 2.

**Table 2:** Comparison of performances of our model with existing method on benchmark dataset.

| Methods | Sensitivity | Specificity | Accuracy | AUROC | MCC |
| --- | --- | --- | --- | --- | --- |
| Sigma70Pred | **97.44** | **97.36** | **97.38** | **0.996** | **0.943** |
| 70ProPred | 92.40 | 96.90 | 95.30 | 0.990 | 0.897 |
| iPro70-FMWin | 83.81 | 95.07 | 91.17 | 0.960 | 0.803 |
| PseZNC | 80.30 | 86.80 | 84.50 | 0.909 | 0.663 |
| Z-Curve | 74.60 | 79.50 | 77.80 | 0.848 | 0.527 |
| IPMD | 82.40 | 90.70 | 87.90 | - | 0.731 |

### Performance comparison on independent dataset

In order to evaluate the method’s robustness and performance, we have also performed testing of our model on the independent dataset of DNA sequences extracted from Regulon DB 10.8, using various existing methods. The results on testing data show that our model is quite robust towards the unseen data and performs well on it. It also implies that our SVM model is significantly free from bias and overfitting on training data. As shown in Table 3, two out of four methods are not able to produce the results and Sigma70Pred outperforms the iPro70-FMWin.

**Table 3:** Performance of existing methods on independent dataset.

| Methods | Sensitivity | Specificity | Accuracy | AUROC | MCC |
| --- | --- | --- | --- | --- | --- |
| Sigma70Pred | **91.45** | **88.56** | **90.41** | **0.95** | **0.79** |
| iPro70-FMWin | 84.12 | 86.67 | 85.04 | 0.92 | 0.69 |
| PseZNC | NW | NW | NW | NW | NW |
| 70ProPred | NW | NW | NW | NW | NW |

### Implementation of model in web server

In order to serve the scientific community, we have also developed the webserver Sigma70Pred by implementing our best model to predict the sigma70 promoters. The web server consists of three modules namely “Predict,” “Scan,” and “Design.” The detailed description of each module is as follows:

### Predict

This module allow users to classify the submitted sequence as sigma70 promoter or non-promoter. There is a restriction of length in tis module as the model is trained on sequences with length 81, hence if the submitted sequence is have length less than 81, ‘A’ will be added as the dummy variable and then the sequence will be classified into one of the class, and if length is greater than 81, only first 81 nucleotide will be considered for prediction. The user can submit sequences in either FASTA or single line format, and can select the desired threshold as SVM score above which the sequence will be classified as sigma70 promoter, otherwise non-promoter. The user can either provide single or multiple sequences, user can also upload the text file containing sequences. The output page display the results in the tabular form, which is downloadable in the csv format.

### Scan

Scan module allow users to scan or identify the sigma70 promoter region in given prokaryote genome. This module does not have any length restriction as in predict module. In this module, overlapping patterns of length 81 will be generated from submitted sequences and then used for prediction. The user can provide single or multiple sequences either in FASTA or in single line format. The user is also allowed to upload the sequence file. The output result will exhibit the overlapping patterns of length 81 with the prediction as promoter or non-promoter. The result is downloadable in the csv format.

### Design

Design module allow users to identify the mutations that can convert the sigma70 promoter into non-promoter or vice-versa. This module also has the restriction of sequence length 81, as it generates all the possible mutants by changing nucleotides at each position and then make the predictions based on the selected threshold. Since, generating all possible mutants is a time and computational expensive process, hence only one sequence is allowed at a time. The output page displays all the possible mutants with its prediction as promoter or non-promoter in tabular form which is downloadable in csv format.

### Standalone

We have also developed python and docker-based standalone package, which is downloadable from URL: https://webs.iiitd.edu.in/raghava/sigma70pred/stand.html. The advantage of this module is that, it is not dependent at the availability of the internet, the user can download these standalones on their local machines and can use all the aforementioned modules. This module also take the input as single or multiple sequences in a file in either FASTA or single line format. The output will be stored in the user defined file in the comma separated value format.

## Conclusions

Sigma70Pred offers a web server and standalone packages to predict the sigma70 promoters using sequence information. This method uses 200 different features, and we assume that our features have more capability to classify sigma70 promoters. Sigma70Pred provide three major modules, such as predict, scan and design. As the application of out method, user can scan the entire prokaryote genome to identify sigma70 promoter, using scan module. By using design module, user can also identify the minimum number of mutations required to exploit the sigma70 promoter region, i.e. either induce or deteriorate the capability of sigma70 promoter. As compared to the existing methods of predicting sigma70 promoters, Sigma70Pred produced commending outcomes. We believe that Sigma70Pred will play an essential role in the area of genomic analysis.

## Acknowledgements

Authors are thankful to funding agencies Department of Biotechnology (DBT), Govt. of India for financial support and fellowships. We are also thankful to Miss Megha Mathur and Anjali Dhall for python scripts to generate features and help in the figures preparation.

## Author contribution

GPSR conceived the idea and supervised the entire project. NS, MZA, and DSP collected and curated the datasets. SP, NS, MZA, and DSP wrote all the in-house scripts, performed the formal analysis and developed the prediction models. SP developed the web interface and standalone. SP and GPSR prepared all the drafts of manuscript. All the authors have given their consent for the final manuscript.

## Reference

1. Paget MS. Bacterial Sigma Factors and Anti-Sigma Factors: Structure, Function and Distribution. Biomolecules 2015; 5:1245–1265.

2. Paget MSB, Helmann JD. The sigma70 family of sigma factors. Genome Biol. 2003; 4:203

3. Bernardo LMD, Johansson LUM, Skarfstad E, et al. sigma54-promoter discrimination and regulation by ppGpp and DksA. J. Biol. Chem. 2009; 284:828–838.

4. Lu C, Xie M, Wendl MC, et al. Patterns and functional implications of rare germline variants across 12 cancer types. Nat. Commun. 2015; 6:10086.

5. Lin H, Li Q-Z. Eukaryotic and prokaryotic promoter prediction using hybrid approach. Theory Biosci. 2011; 130:91–100.

6. Gama-Castro S, Salgado H, Santos-Zavaleta A, et al. RegulonDB version 9.0: high-level integration of gene regulation, coexpression, motif clustering and beyond. Nucleic Acids Res. 2016; 44:D133–43.

7. Song K. Recognition of prokaryotic promoters based on a novel variable-window Z-curve method. Nucleic Acids Res. 2012; 40:963–971.

8. Lin H, Liang Z-Y, Tang H, et al. Identifying Sigma70 Promoters with Novel Pseudo Nucleotide Composition. IEEE/ACM Trans. Comput. Biol. Bioinforma. 2019; 16:1316–1321.

9. He W, Jia C, Duan Y, et al. 70ProPred: a predictor for discovering sigma70 promoters based on combining multiple features. BMC Syst. Biol. 2018; 12:44.

10. Rahman MS, Aktar U, Jani MR, et al. iPro70-FMWin: identifying Sigma70 promoters using multiple windowing and minimal features. Mol. Genet. Genomics 2019; 294:69–84.

11. Friedel M, Nikolajewa S, Suhnel J, et al. DiProDB: a database for dinucleotide properties. Nucleic Acids Res. 2009; 37:D37–40.

12. Chen W, Zhang X, Brooker J, et al. PseKNC-General: a cross-platform package for generating various modes of pseudo nucleotide compositions. Bioinformatics 2015; 31:119– 120.

13. Liu B, Zhang D, Xu R, et al. Combining evolutionary information extracted from frequency profiles with sequence-based kernels for protein remote homology detection. Bioinformatics 2014; 30:472–479.

14. Pedregosa F, Varoquaux G, Gramfort A, et al. Scikit-learn: Machine Learning in Python. J. Mach. Learn. Res. 2011; 12:2825–2830.

15. Patiyal S, Agrawal P, Kumar V, et al. NAGbinder: An approach for identifying N□acetylglucosamine interacting residues of a protein from its primary sequence. Protein Sci. 2020; 29:201–210.

16. Sachs MC. plotROC: A Tool for Plotting ROC Curves. J. Stat. Softw. 2017; 79.

