## Supplementary file for "Sigma70Pred: A highly accurate method for predicting sigma70 promoter in prokaryotic genome"

**1. Feature Generation**

In order to develop the prediction model with the ability to classify sigma70 promoters, we have generated wide range of features. The description of each feature is as follows:

1.1 **Position-Specific Trinucleotide Propensity (PSTNPP)**

Position-Specific Trinucleotide Propensity [1] is based on the idea that position-specific values of nucleotides and their k-mers possess special properties and can be utilized in the development of Machine Learning models for the tasks of promoter identification. Many studies have proposed an idea to generate the composition of trimers in a nucleotide sequence for a particular position. The steps to generate these values are as follows:

We have considered nucleotide sequences of length = 81 for our tasks. Thus, a single sample sequence would be represented as,

X = N_1_N_2_N_3_N_4_……...N_81._ (Single Sequence)

*Where, N_i_ can be A, C, G, T and i represent the position of a nucleotide.*

A trimer is the combination of three nucleotides and if we count sequentially we will get 79 trimers for a sequence with length 81. If a sequence has length L then L-2 is the number of trimers the sequence can have. There are 4 different nucleotides (A, C, G, T), and with the combination of three nucleotides, we can generate 4^3^ = 64 different trimers such as AAA, AAC, AAG, etc. Hence, when we generate 79 position-specific compositions for 64 different trimers, we get a matrix of dimension 64 X 79. The matrix can be represented as:

|  | **Z_1,1_** | **Z_1,2_** | **…** | **Z_1,79_** |
| --- | --- | --- | --- | --- |
|  | **Z_2,1_** | **Z_2,2_** | **…** | **Z_2,79_** |
| **Z =** | **.** | **.** | **…** | **.** |
|  | **.** | **.** | **…** | **.** |
|  | **Z_63,1_** | **Z_63,2_** | **…** | **Z_63,79_** |
|  | **Z_64,1_** | **Z_64,2_** | **…** | **Z_64,79_** |

*Where,*

Z_i,j_ = Q**_+(i,j)_** – Q**_-(i,j)_**

Q**_+(i,j)_** stands for the count of i^th^ trimer at position j in positive sequences.

Q**_-(i,j)_** stands for the count of i^th^ trimer at position j in negative sequences.

Using the matrix Z, we can convert each nucleotide sequence into a vector of length 79 where each value in a vector denotes the difference in the counts of the trimer in the positive and negative sequences at a particular position under consideration. The idea is illustrated below:

S = ATGCAGT…………. ACC (Sample Sequence of length 81)

Here, the first trimer i.e at i=1 is *ATG*. In the corresponding feature vector form, the ATG will be replaced by the value of ATG in a row for ‘ATG’ in the matrix ‘Z’ and the column would be 1, i.e it denotes the difference of count of trimer ‘ATG’ in positive and negative sigma70 sequences at position 1.

Similarly, we would repeat the procedure for the second and succeeding trimers incrementing the ‘i’ (column number) at each step, whereas row number is defined by the trimer itself. Finally, we will get the vector feature V as:

V = [ v_1_, v_2_, v_3_, ........., v_79_] (Vector form of a sequence)

v_1_, v_2_, and so on denotes the position-based composition values taken from matrix Z.

To derive row number for each trimer the following table is used where the sequence of nucleotide is taken as *“ACGT”.* The trimer combination derived from this sequence would give the corresponding row number to each trimer as follows:

| **Trimer** | **Row Number** |
| --- | --- |
| AAA | 1 |
| AAC | 2 |
| AAG | 3 |
| --- | --- |
| --- | --- |
| TTT | 64 |

**1.2 Electron-Ion interaction pseudopotentials of trinucleotide (EIIIP)**

EIIIP [1] is a feature generation method for DNA sequence where we use the Electron-Ion potential values of each nucleotide (A, C, G, T) to create a feature vector of dimension *64* using each trinucleotide present in the DNA sample sequence. The general form of each feature vector in the EIIP method is:

***V = [ EIIP_AAA_^.^N_AAA_, EIIP_AAC_^.^N_AAC_, ............, EIIP_TTT_^.^N_TTT_]***

*where,*

EIIP_xyz_ denotes the EIIP value of trinucleotide XYZ.

N_xyz_ denotes the Normalized Frequency of trinucleotide XYZ in a given sequence.

Further, we could derive EIIP value as:

EIIP_xyz_ = E_x_ + E_y_ + E_y_

*where,*

E_x_ denotes the Electron-Ion Values for the Nucleotide X **∈** (A, C, G, T).

Refer to the following table to see the EIIP Value of Nucleotides:

| **Nucleotide** | **EIIP** |
| --- | --- |
| **A** | **0.1260** |
| **C** | **0.1340** |
| **G** | **0.0806** |
| **T** | **0.1335** |

**1.3 Dimer Count**

The dimer is the two-length combination of nucleotides ( A, C, G, T ) present in a DNA sequence [2]. The dimer could be represented as:

*D_k_ = N_i_N_j_*

*where,*

i and j represent nucleotides present at different positions in the sequences.

D_k_ represents the k^th^ dimer formed by combinations of nucleotides.

For each N_i_ and N_j_, we have 4 choices for nucleotide. Thus, the total number of unique dimers that can be formed are 4 X 4 = 16 *(AA, AC, ..., TT)*. We have calculated the composition of each dimer in a sequence. The counts of the dimer are calculated with different gaps in between two nucleotides of the dimer. The idea is illustrated below:

X = N_1_N_2_N_3_N_4_N_5_…...N_81_ (sample sequence of length 81)

N represents the i^th^ nucleotide.

*A dimer with no gap would be formed by*-

N_1_	N_2_	N_3_	N_4_	N_5_…… N_81_

The corresponding dimers are - N_1_N_2_, N_2_N_3,_ and so on.

*A dimer with a gap of 1 would be formed by*-

N_1_	N_2_	N_3_	N_4_	N_5_…… N_81_

The corresponding dimers are - N_1_N_3_, N_2_N_4_, and so on.

Similarly, we could form dimers with gaps 2,3,4, and so on. We have considered dimers without gap and with gaps of 1, 2, 3, 4, and 5 as features for the classifier.

1.4 **Trimer Count**

The trimer is the three-length combination of nucleotides (A, C, G, T) present in a DNA sequence [2]. The dimer could be represented as:

*D_k_ = N_x_N_y_N_z_*

*where,*

X, Y, and Z represent nucleotides present at different positions in the sequences.

D_k_ represents the k^th^ trimer formed by combinations of nucleotides.

For each N we have 4 choices of nucleotides. Thus the total number of unique trimers that can be formed are 4X4X4 = 64 (AAA, AAC,..., TTT).

We have calculated different forms of trimers (gap, no-gap) and their composition in each DNA sequence. The idea is illustrated below:

Considering the sequence (as in the previous dimer case), *a trimer with no-gap would be represented by-*

The corresponding trimers are- N_1_N_2_N_3_, N_2_N_3_N_4_, and so on.

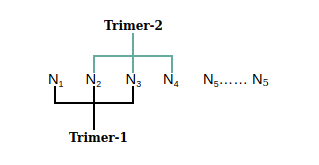

Trimer with Left Gap –

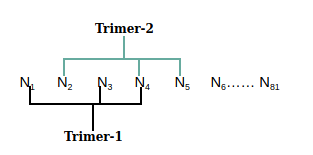
Taking Gap = 1

The corresponding trimers are, N_1_N_3_N_4_, N_2_N_4_N_5_, and so on.

*Trimer with Right Gap-*

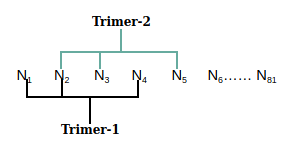

The above figure represents the trimers generated after *taking gap=1*. Similarly, we can take different gaps (2,3,4 and so on) and can generate more trimers. In our feature generation, we have taken gap values as - 1, 2, 3, 7, 8, 9, 10, 15, 16, 17 to generate trimers. (Both for left gap and right gap).

1.5 **Motif Counts**

These are the special patterns in the DNA sequence that may repeat their occurrence and are responsible for biological activities related to DNA [2]. Some of the patterns occurring in DNA sequences are useful for identifying promoter regions. We have used the count of some of these patterns for our feature construction phase. We have considered the following 9 patterns in our approach:

TTGAC, TATAAT, TTATATA, TTGACA, AACGAT, ACAGTT, AGGAGG, TAAAAT and TTGATT.

1.6 **GC and AT Skew**

In order to capture the relative-frequency of DNA bases in a given sequence, we have used GC Skew [2]*.* The GC skewness is represented as:

*GC skew = (G − C) / (G + C)*

*here,*

G and C stand for the count of nucleotides Guanine and Cytosine, respectively.

Similarly, we have also evaluated the AT Skew in each DNA sequence as:

*AT skew = (A − T) / (A + T)*

A and T stand for the count of nucleotides Adenine and Thymine, respectively.

1.7 **Nucleotide Composition Features:**

The correlation-based features of DNA sequences are also explored. Correlation is basically a relation between properties. There are two types of correlations, If the property is related to its own then it is termed **auto-correlation** and if there exists some correlation between two properties then it is termed s **cross-correlation**. Correlation-based features basically convert the different length DNA sequences into fixed-size vectors.

The following are some of the features of this type included in the system:

**1.7.1 Dinucleotide autocorrelation (DAC):** The correlation between the same physicochemical indices for two dinucleotides separated by a distance of lag is measured by the function [3].

$$\boldsymbol{DAC}\left( \boldsymbol{\mu}\boldsymbol{,lag} \right)\boldsymbol{=}\sum_{\boldsymbol{i=1}}^{\boldsymbol{L-lag-1}} \boldsymbol{P}_{\boldsymbol{\mu}}\left( \boldsymbol{N}_{\boldsymbol{i}}\boldsymbol{N}_{\boldsymbol{i+1}} \right)\boldsymbol{-}{\bar{\boldsymbol{P}}}_{\boldsymbol{\mu}}\boldsymbol{)/(L-lag-1)}$$

*here,*

$${\bar{\boldsymbol{P}}}_{\boldsymbol{\mu}}\boldsymbol{=}\sum_{\boldsymbol{j=1}}^{\boldsymbol{L-1}} \boldsymbol{P}_{\boldsymbol{\mu}}\boldsymbol{(}\boldsymbol{N}_{\boldsymbol{i}}\boldsymbol{N}_{\boldsymbol{i+1}}\boldsymbol{)/(L-1)}$$

1.7.1 **Dinucleotide cross-correlation (DCC):** The correlation between two different physicochemical indices for two different dinucleotides separated by a distance of lag is measured by the function [3].

$$\boldsymbol{DCC}\left( \boldsymbol{\mu}_{\boldsymbol{1}}\boldsymbol{,}\boldsymbol{\mu}_{\boldsymbol{2}}\boldsymbol{,lag} \right)\boldsymbol{=}\sum_{\boldsymbol{i=1}}^{\boldsymbol{L-lag-1}} \frac{\boldsymbol{P}_{\boldsymbol{\mu}_{\boldsymbol{1}}}\left( \boldsymbol{N}_{\boldsymbol{i}}\boldsymbol{N}_{\boldsymbol{i+1}} \right)\boldsymbol{-}{\bar{\boldsymbol{P}}}_{\boldsymbol{\mu}_{\boldsymbol{1}}}\boldsymbol{)(}\boldsymbol{P}_{\boldsymbol{\mu}_{\boldsymbol{2}}}\left( \boldsymbol{N}_{\boldsymbol{i+lag}}\boldsymbol{N}_{\boldsymbol{i+lag+1}} \right)\boldsymbol{-}{\bar{\boldsymbol{P}}}_{\boldsymbol{\mu}_{\boldsymbol{2}}}\boldsymbol{)}}{\boldsymbol{(L-lag-1)}}$$

*here,*

$${\bar{\boldsymbol{P}}}_{\boldsymbol{\mu}}\boldsymbol{=}\sum_{\boldsymbol{j=1}}^{\boldsymbol{L-1}} \boldsymbol{P}_{\boldsymbol{i}}\boldsymbol{(}\boldsymbol{N}_{\boldsymbol{j}}\boldsymbol{N}_{\boldsymbol{j+1}}\boldsymbol{)/(L-1)}$$

1.7.2 **Dinucleotide Auto Cross-Correlation (DACC):** This function calculates the dinucleotide auto cross-correlation i.e. it is a mixture of DAC and DCC [3].
1.7.3 **Moran Auto-Correlation (MAC):** MAC stands for Moran auto-correlation. It computes the correlation of the same property between two nucleotides (kmer -1,2 or 3) separated by a distance of lag [4].

$$\boldsymbol{MAC}\left( \boldsymbol{\mu}\boldsymbol{,k,lag} \right)\boldsymbol{=}\frac{\left[ \frac{\boldsymbol{1}}{\boldsymbol{L-lag-k+1}} \right]\sum_{\boldsymbol{i=1}}^{\boldsymbol{L-lag-k+1}} \boldsymbol{(P}_{\boldsymbol{\mu}}\left( \boldsymbol{x}_{\boldsymbol{i}} \right)\boldsymbol{-}{\bar{\boldsymbol{P}}}_{\boldsymbol{\mu}}\boldsymbol{(x))(}\boldsymbol{P}_{\boldsymbol{\mu}}\left( \boldsymbol{x}_{\boldsymbol{i+log}} \right)\boldsymbol{-}{\bar{\boldsymbol{P}}}_{\boldsymbol{\mu}}\boldsymbol{(x))}}{{\boldsymbol{(}\frac{\boldsymbol{1}}{\boldsymbol{L-k+1}}\sum_{\boldsymbol{i=1}}^{\boldsymbol{L-k+1}} \boldsymbol{(}\boldsymbol{P}_{\boldsymbol{\mu}}\left( \boldsymbol{x}_{\boldsymbol{i}} \right)\boldsymbol{-}{\bar{\boldsymbol{P}}}_{\boldsymbol{\mu}}\left( \boldsymbol{x} \right)\boldsymbol{)}}^{\boldsymbol{2}}}$$

1.7.4 **Normalized Moreau-Broto autocorrelation (NMBAC):** It computes the correlation of the same property between two nucleotides (kmer-1,2 or 3) separated by a distance of lag [4].

$$\boldsymbol{NMBAC}\left( \boldsymbol{\mu}\boldsymbol{,k,lag} \right)\boldsymbol{=}\frac{{\sum_{\boldsymbol{i=1}}^{\boldsymbol{L-lag-k+1}} \boldsymbol{(}\boldsymbol{P}_{\boldsymbol{\mu}}\left( \boldsymbol{x}_{\boldsymbol{i}} \right)\boldsymbol{\times}\boldsymbol{P}_{\boldsymbol{\mu}}\left( \boldsymbol{x}_{\boldsymbol{i+lag}} \right)\boldsymbol{)}}^{\boldsymbol{2}}}{\boldsymbol{L-k-lag+1}}$$

1.7.4.1 **Parallel Correlation Pseudo Tri-Nucleotide Composition (PC_PTNC):** This function computes parallel correlation pseudo Trinucleotide composition [5].

At the end of the feature generation phase, we have calculated a feature vector of length 8465 for each DNA sequence of 81 bp.

**2. Feature Scaling**

We have used the Min-Max scaler from the scikit-learn library [6] to scale down the values of the feature we have constructed. This is done in order to bring the values of features to the same range (usually 0 and 1).

Mathematically, min-max scaling using scikit-learn can be described as:

$$Q_{std}=\frac{(Q-Q_{min})}{(Q_{max}-Q_{\min})}$$

$$Q_{scaled}= Q_{std}*\left( maximum-minimum \right)+minimum$$

*here,*

Q stands for the value of a particular feature.

Q_min_ is the minimum value of the feature X in the given dataset.

Q_max_ is the maximum value of the feature X in the given dataset

maximum and minimum define the feature range.

**3. Feature Selection**

We have used *Recursive Feature Elimination* (RFE) [6] for the feature selection. In this technique, we eliminate some of the features from our dataset and construct the model using the base estimator function on a new set of features, and performance is evaluated on the basis of certain metrics. This process is repeated until we are left with the desired number of features. The features are weighted as per the evaluation criteria in order to remove or consider a particular feature. We have used the scikit-learn module for the RFE evaluation on our training dataset.

In order to find the optimal number of features, we have used the RFE multiple times with the different number of final features, on our dataset. We have used Logistic Regression as the base classifier in the RFE. The final parameters to the RFE are estimator=LogisticRegression(), n_features_to_select=200, step=10. We finally reduced the dimension of our 8465 feature vector to 200 features.

**Reference:**

[1]	W. He, C. Jia, Y. Duan, Q. Zou, *BMC systems biology* **2018**, *12*, 44-44.

[2]	M. S. Rahman, U. Aktar, M. R. Jani, S. Shatabda, *Molecular genetics and genomics : MGG* **2019**, *294*, 69-84.

[3]	M. Friedel, S. Nikolajewa, J. Suhnel, T. Wilhelm, *Nucleic Acids Res* **2009**, *37*, D37-40.

[4]	W. Chen, X. Zhang, J. Brooker, H. Lin, L. Zhang, K. C. Chou, *Bioinformatics* **2015**, *31*, 119-120.

[5]	B. Liu, D. Zhang, R. Xu, J. Xu, X. Wang, Q. Chen, Q. Dong, K. C. Chou, *Bioinformatics* **2014**, *30*, 472-479.

[6]	F. Pedregosa, G. Varoquaux, A. Gramfort, V. Michel, B. Thirion, O. Grisel, M. Blondel, P. Prettenhofer, R. Weiss, V. Dubourg, J. Vanderplas, A. Passos, D. Cournapeau, M. Brucher, M. Perrot, E. Duchesnay, *Journal of Machine Learning Research* **2011**, *12*, 2825-2830.
